# Reconstruction of *Escherichia coli* ancient diversification by layered phylogenomics and polymorphism fingerprinting

**DOI:** 10.1101/486399

**Authors:** José M González-Alba, Fernando Baquero, Rafael Cantón, Juan Carlos Galán

**Author notes:** **Corresponding author: E-mail** (JCG).

## Abstract

The rapidly increasing availability of whole genomes provides the opportunity to reach an updated comprehensive view of bacterial evolution. The staggered diversification of evolutionary processes, based on the combined strategy of layered phylogenomics and polymorphism fingerprinting, give a new perspective in phylogenetic reconstructions. Layered phylogenomics is based on the assignation of genes according to five different evolutionary layers: minimal genome, genus-core genome, species-core genome, phylogroup-core genome and phylogroup-flexible genome. Polymorphism fingerprinting is based on the detection of conserved positions in each phylogenetic group but differing from those of their hypothetical ancestors. This approach was applied to *Escherichia coli* because there are unresolved evolutionary questions, although has been highly studied. Phylogenetic analysis based on 6,220 full genomes, identified three *E. coli* root lineages, defined as D, EB1A and FGB2. A new phylogroup, called G was detected near to phylogroup B2. The closest phylogroup to ancestral *E. coli* was phylogroup D, whereas E and F were the closest ones in their respective lineages; moreover, A and B2 were the most distant phylogroups in EB1A and FGB2 respectively. We suspect that EB1A and FGB2 lineages represent different adaptive strategies. In the deepest branch of EB1A lineage, the number of accumulated mutations was lower than in recent branches, whereas in FGB2 lineage the opposite occurred. The FGB2 lineage was enriched in genes related to host colonization-pathogenicity and toxin-antitoxin systems (such as *hip*A), whereas B1A sub-lineage acquired functions related to uptake and metabolism of carbohydrates (such as *bgl, mng* or *xly*E). This new combined strategy shows a detailed staggered evolutionary reconstruction, which help us to understand the deepest events and the selection forces have driven *E. coli* diversification. This approach could add resolution in the reconstruction of the evolutionary trajectories of other microorganisms.

**Author summary:** Phylogeny based on whole genome provides the opportunity to study the history of eco-adaptive diversification of any bacterial taxon. Different strategies have been proposed for knowing the evolutionary trajectories in some species, such as *Escherichia coli*; however, these analyses were based on a limited number of sequences, and sometimes the evolutionary reconstructions reached clashed positions, especially in the ancestral inferences. For adding resolution in evolutionary reconstructions, we propose a combination of approaches, such as layered phylogenomics based on the use of different set of genes corresponding to the successive evolutionary steps, and polymorphism fingerprinting which detects hallmarks of the ancient mutations. We propose to use *E. coli* because it is paradigmatic example of the evolutionary inconsistences despite being a microorganism with enough evolutionary analysis. Three ancestral lineages were established with this strategy and the staggered reconstruction about the origin and diversification of *E. coli* phylogroups was inferred. Moreover, in the context of this study, a new *E. coli* phylogroup was defined. The main lineages represent different adaptive strategies, one lineage gained genes involved in pathogenicity, and another one acquired genes allowing the obtainment of energy from different sources.

## Introduction

Since the first description by Theodore Escherich described of *Escherichia coli* in 1885, several generations of researchers have been fascinated by this organism. *E. coli* has been extensively used as a model to understand bacterial adaptability [1, 2]. The population diversity of *E. coli* was initially recognized in four main phylogroups (A, B1, B2 and D) [3]. In the following years, the increasing number of available sequences allowed the identification of three new phylogroups (C, E, and F) and five cryptic clades, revealing that the population structure of *E. coli* was more complex than initially suggested [4]. When the first whole *E. coli* genome was sequenced in 1997, a new possibility in the comparative genomic field was perceived for this microorganism [5]. The growing availability of a large amount of whole *E. coli* genomes provided an unprecedented level of discrimination and the opportunity to perform solid evolutionary reconstructions [6]. Traditionally, the bacterial genomes have been distinguished in a core genes pool encoding the basic cellular functions, and a flexible genes pool conferring strain-, pathotype- or ecotypes-specific characteristics which allow adaptation to special conditions [7]. For instance, from the first available studies based on a limited number of sequences, ranging from 20-61 genomes [8, 9], until the most recent ones using fewer than 250 genomes [10, 11], several discrepancies particularly regarding the origin and ancestral position of the different lineages are still unresolved. Some groups considered phylogroup B2 to be the most ancestral *E. coli* phylogroup [8, 2, 10], whereas other studies proposed phylogroup D was in this ancestral position [12]. On the other hand, it was also suggested that two D sub-lineages could be the origin of two main evolutionary trajectories leading to lineages A/B1/C/E and B2/F [13, 6, 14]. Other researchers suggested that phylogroup B1 was the origin of E and A phylogroups [1] or proposed a paraphyletic origin for phylogroup A [15]. Other works questioned the differentiation of phylogroup C [12, 16].

To contribute to answer these unresolved evolutionary questions by adding resolution in the evolutionary reconstructions, a new strategy was proposed to elucidate the successive steps in the *E. coli* diversification. Our strategy was a combination of two approaches. One of them, coined “layered phylogenomics” (LP) is based on stratified phylogenetic analysis of genes representing successive evolutionary steps. The layers are divided in minimal genome, genus-core genome species-core genome, phylogroup-core genome and phylogroup-flexible genome (Fig 1). The LP approach was complemented with the “polymorphism fingerprinting” (PF) approach, based on the identification of the conserved positions in each phylogenetic group, that are variable with respect to their hypothetical ancestor. This strategy could allow a visual representation of the staggered diversification processes of *E. coli*.

**Fig 1.**
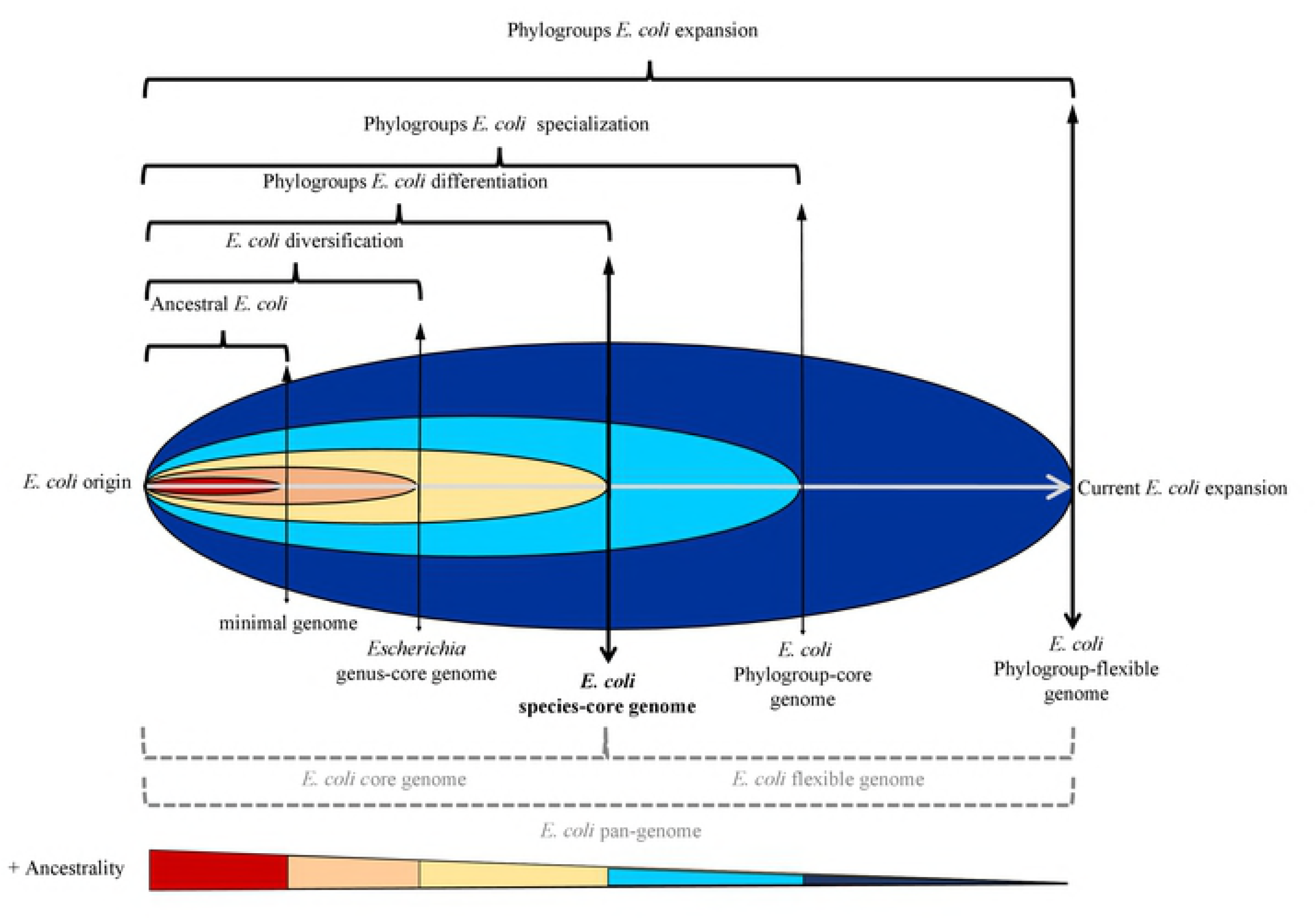
Proposal of framework for the evolutionary reconstruction of *E. coli*. The bacterial DNA classically allocated in core or flexible genomes (thick vertical lines) were subdivided in order to obtain an evolutionary gradient from the most ancestral genes (core genome) to the recently acquired (flexible genome). Different layers of analysis, reflecting the taxonomic units genetically established (bacteria, genus, species, phylogroup) and remaining genes were considered. The most ancestral set of genes corresponds to those genes identified as minimal genome (red), representing genes present in all bacteria and probably they are essential genes. *Escherichia* genus-core genome (orange) corresponds to the genes implicated in the *Escherichia* diversification prior to the formation of *E. coli* species. *E. coli* species-core genome (yellow), represents the period between the emergence of the species until the start of *E. coli* specialization. Phylogroup-core genome (light blue) represents the specialization and phylogroup-flexible genome including the remaining genes (dark blue), to reach the current limit *E. coli* expansion.

## Results

### Defining the framework for the evolutionary reconstruction of *E. coli*

At the time of starting this work, the number of available *E. coli* sequences in the genome database from NCBI was 6,266 genomes. To ascertain if all *E. coli* genomes were correctly identified, the core genome of *Escherichia* genus was established in 189 genes shared by all members. The phylogenetic tree constructed with the concatenated sequence of these genes revealed that 40 genomes were wrongly classified as *E. coli* mainly belonged to cryptic clade I, the closest related lineage to *E. coli* (S1 Fig). Moreover, another six genomes were also excluded because their poor sequencing. Once the remaining 6,220 genomes were confirmed, *E. coli* core genome was established in 1,027 genes. A phylogenetic tree was constructed with these genes and was used as the reference phylogeny. This tree confirmed most of the previously known *E. coli* phylogroups, but we were unable to unequivocally separate phylogroup C from B1. On the contrary, a new phylogroup was found, which we proposed to designate as phylogroup G, following the pre-established denomination (Fig 2A). The estimation of evolutionary divergences over sequences pairs between phylogroups reinforced the identification of phylogroup G (Fig 2B). This phylogroup is a monophyletic clade with low diversity, located next to phylogroup B2. Two *E. coli* genomes (KTE146 and EPEC-503225) were located in an intermediate position between the node of *Escherichia* cryptic clade I and the origin of the *E. coli* diversification. Nowadays, these sequences could be used as better candidates than cryptic clade I in the ancestral reconstruction of *E. coli* diversification as the evolutionary distance between cryptic clade I and *E. coli* origin is too large to be considered as the most recent ancestor.

**Fig 2.**
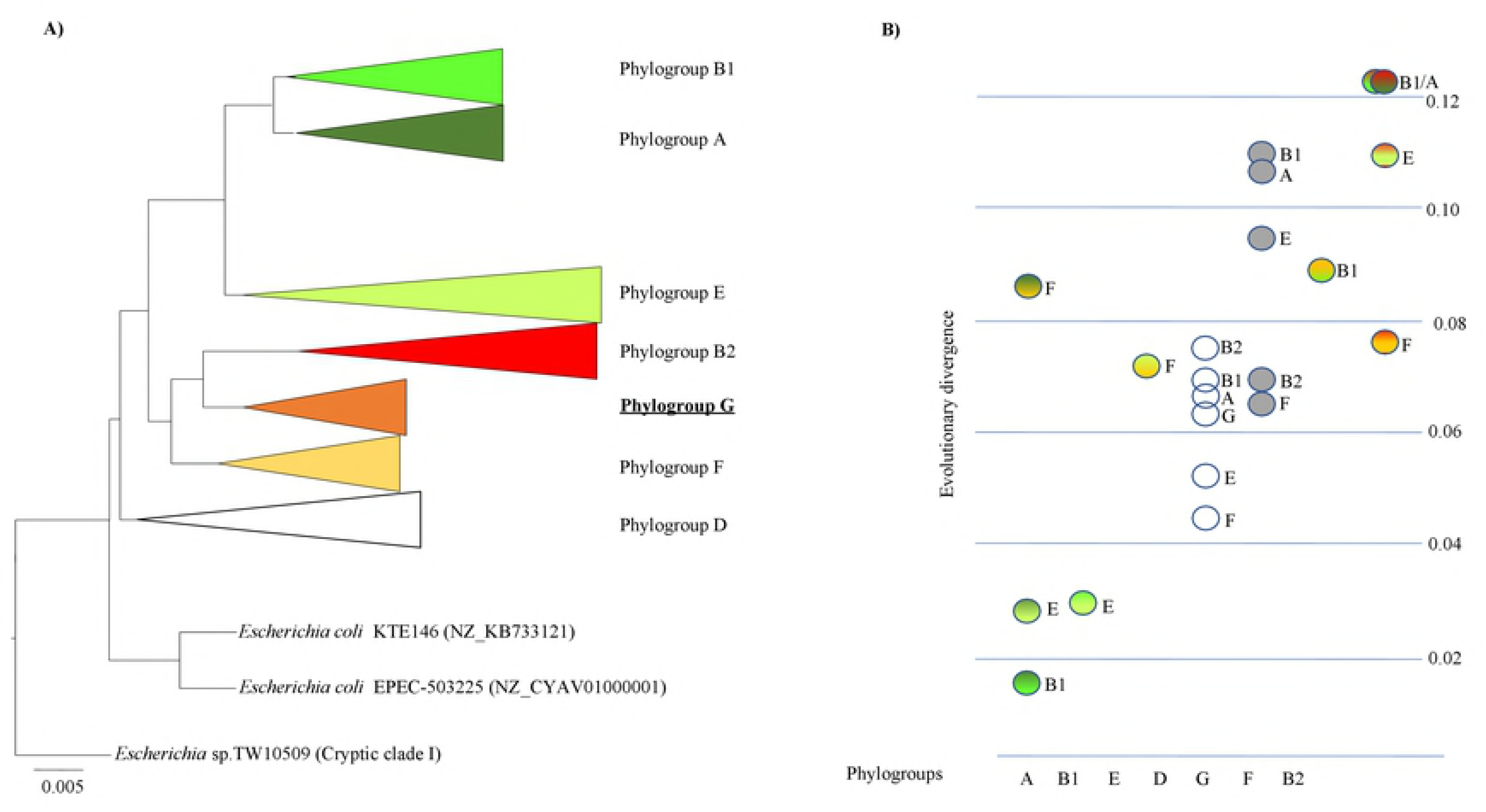
*Escherichia coli* species phylogenetic reconstruction and evolutionary divergence among the phylogroups. **A**) ***E. coli* species core phylogeny**. Phylogenetic reconstruction using Maximum-likelihood (GTR+I+Γ, SH ≥99%) with the concatenate of 1,027 genes (1,046,053 nt) present in the 100% of sequenced *E. coli* strains. All concatenate with less than 95% site coverage were eliminated. The established phylogroup C could not be distinguished in phylogroup B1. Cryptic clade I was included as outgroup in the reconstruction. **B**) **Estimates of average evolutionary divergence over all sequence pairs between groups**. The number of base substitutions per site from averaging all sequence pairs between groups pairs using the GTR+I+ Γ model are shown. The evolutionary distance, indicated as circles with different colors and a character, correspond to the distance between the phylogroup in X-axis and the phylogroups indicated for the character next to the circle. The grey circles show the distance of the new identified phylogroup G. The white circles correspond to the evolutionary divergence of phylogroup D. The circles with two colors correspond to the comparison between the other phylogroups using the same colors as Fig 2A

The *E. coli* core genome phylogeny also suggested three root lineages. They were denominated as EB1A (including the E, B1 and A phylogroups), FGB2 (including F, G, and B2 phylogroups) and D (including phylogroup D). Among the phylogroups allocated in the lineage EB1A, 859 sequences were identified as phylogroup E (average chromosomal size 5,364.150), 1,995 as phylogroup B1 (average chromosomal size 5,197.510) and 1,296 as phylogroup A (average chromosomal size 4,977.757). Meanwhile, in lineage FGB2 the distribution was: 124 sequences corresponded to phylogroup F (average chromosomal size 5,321.950), 55 to phylogroup G (average chromosomal size 5,245.213) and 1,455 to phylogroup B2 (average chromosomal size 5,138.164). Finally, 424 sequences were attributed to phylogroup D (average chromosomal size 5,252.449) and 10 sequences could not be allocated to any known phylogroup (note that the number of genomes per phylogroup does not necessarily reflect the *E. coli* population distribution). A representation of chromosomal sizes is shown in S2 Fig.

### LP-PF strategy yield detailed evolutionary reconstruction of the origin of *E. coli* phylogroups

Now, we envisaged elucidating the evolutionary steps leading to such phylogeny using the LP approach, detailed in the Material and Methods section and S3 Fig.

In first layer corresponding to known as minimal genome, only 51 among the previously described genes [17, 18] were found in all *E. coli* genomes. In *Escherichia* genus-core genome (second layer), 189 genes were found; however, from this number we subtracted the genes from minimal genome, in order to reconstruct the corresponding phylogenetic tree based on 138 genes of genus-core genome. In the third layer, the *E. coli* species-core genome was reconstructed using 838 genes, after excluding the 189 from *Escherichia* genus-core genome. The LP approach revealed identical topology with the three set of genes used (S4A Fig) suggesting that this approach was still insufficient to infer the first steps in the differentiation and diversification of *E. coli*. Consequently, we conceived the possibility of complementing recognizing patterns of single nucleotide polymorphisms (SNPs) accumulated along the *E. coli* evolutionary steps used in the above section. These SNPs could be used as high-support markers (fingerprinting) in the ancestral reconstruction of phylogenetic groups, which we called the phylogroup “polymorphism fingerprinting” (PF) approach, adding resolution to the reconstruction observed with only LP approach. As expected, the percentages of invariable positions were higher among the genes belonging to minimal genome, 81% (42,495 invariable positions/52,404 total positions), followed by 78% (150,108/191,766) among the genes classified as *Escherichia* genus-core and 75% (601,412/801,883) in *E. coli* species-core genome. The phylogroup- or lineage-specific changes present in all genomes were identified. A total number of 14 (3‰), 88 (5‰) or 490 (8‰) mutations were defined as specific in the minimal genome, *Escherichia* genus-core genome and *E. coli* species-core genome respectively. Subsequently the numbers of specific changes were overprinted in the corresponding branches of the phylogenetic trees (S4B Fig).

The combined strategy (LP-PF) was suggestive of a more detailed evolutionary scenario, from the deepest branches to reaching the latest events in the differentiation processes of the classic *E. coli* phylogroups. Therefore, we can propose a staggered evolutionary scenario in Fig 3. Lineage D, with phylogroup D as unique member showed fewer changes with respect to known most recent common ancestor (MRCA) than other lineages, and then we assumed that it was the last phylogroup separated from the ancestral genome and consequently the lineage more closely related to *E. coli* origin. This LP-PF strategy also allowed us to infer the successive diversification steps in EB1A and FGB2 lineages. In FGB2 lineage, we were able to identify phylogroup F as the last group separated from FGB2 root but not to identify which one was the first diverging phylogroup (B or G). Reconstruction of EB1A lineage only allowed us to suggest the appearance of the EB1A lineage as a non-ancient step and the subsequent separation of the B1A sub-lineage (Fig 3).

**Fig 3.**
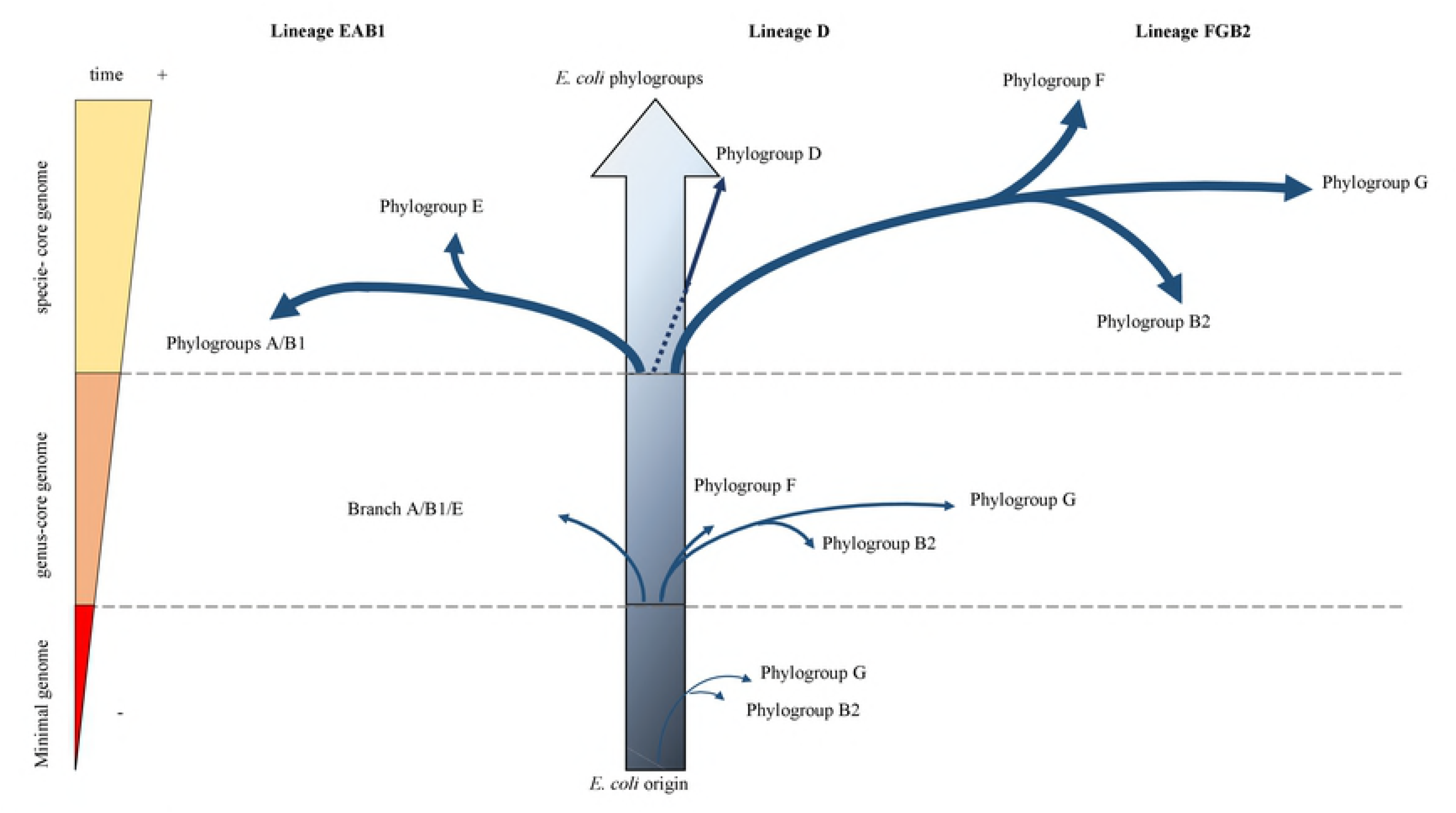
Proposed evolutionary scenario in the diversification of *E. coli* based on results obtained with LP-PF strategy. The branches reflect the accumulated mutations, but their lengths are not proportional to the observed distance.

To reinforce this evolutionary scenario, we explore the gain and loss of ancient genes reconstructing the hypothetical ancient *E. coli* core genome based on the phylogroup-core genomes, the fourth evolutionary layer in our model [19]. The gene content of phylogroups-core genomes ranged from 741 to 2,715 genes, corresponding to phylogroups A and G respectively, once the 1,027 genes corresponding to *E. coli* core genome were excluded. A set of 2,052 genes constituting this ancient genome was searched in all individual genomes of each phylogroup. These data permitted calculation of the percentage of genomes in each phylogroup carrying 95-99% of ancient genes. When the threshold of ancient genes was 95%, no differences among phylogroups were detectable; however, the step-wise increase of this threshold towards 99% progressively revealed differences among them (Fig 4). Consistently with the previous analysis, phylogroup D maintained the highest percentage of strains sharing 99% of ancient genes, supporting that this phylogroup was the ancestral one. Now, phylogroup B2 was the first in FGB2 lineage to be separated from the hypothetical ancestral genome, and phylogroup F was the last one, confirming the previous results. Inside the EB1A lineage, phylogroup A was the first differentiated member, while phylogroup was E was the last one separated from the ancestral genome. Moreover, EPEC-503225 and KTE146 strains carried 99% of the ancestral genes, supporting our proposal that these strains could represent the best-to-the-present known close ancestors of *E. coli*.

**Fig 4.**
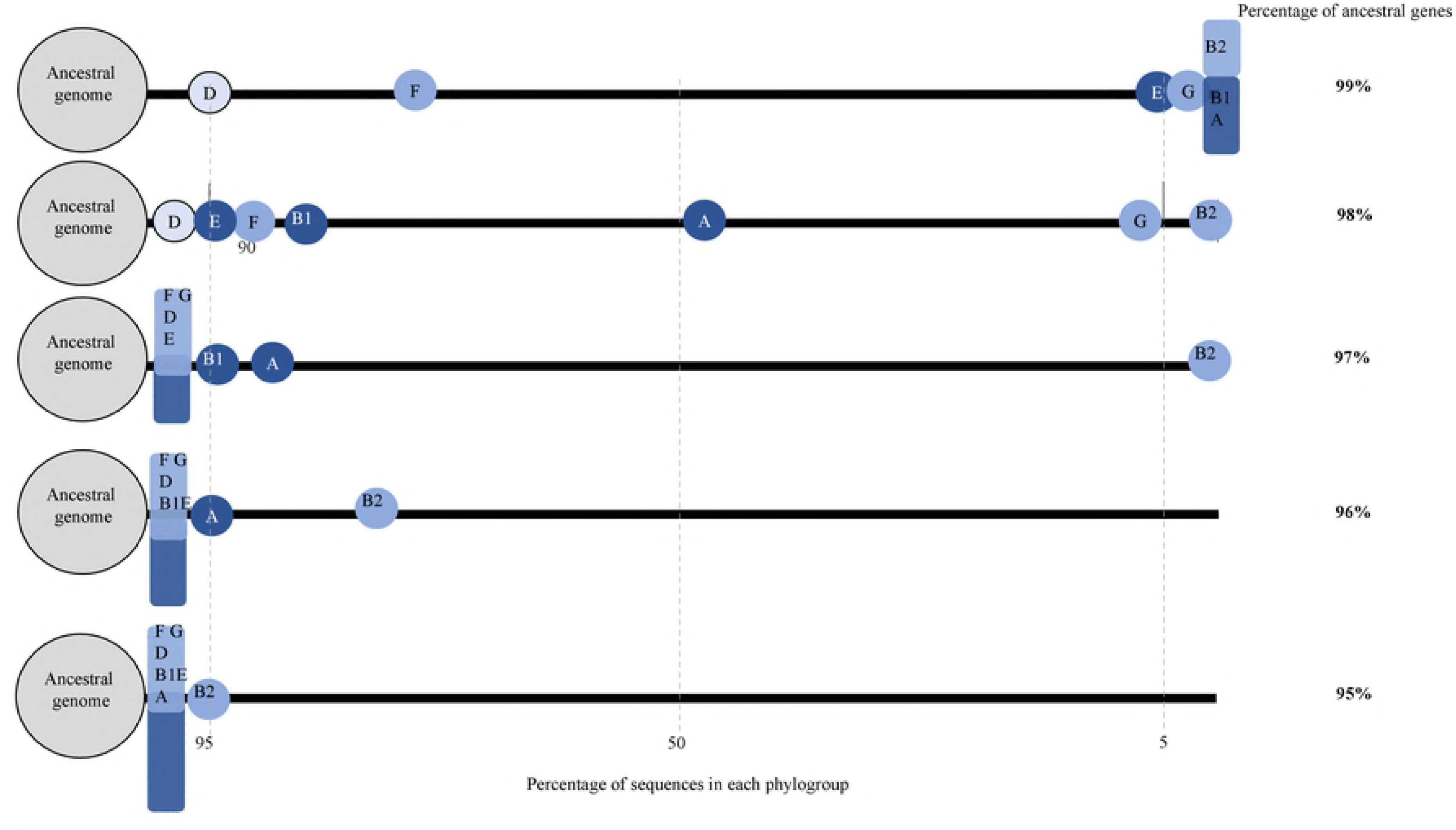
Representation of the presence of ancestral genes in each phylogroup. The percentages of strains carrying 95-99% of genes identified as ancient genome are represented using the MGRA program (http://mgra.cblab.org). In the right column the percentage of genes from hypothetical ancient genome conserved in each phylogroup is presented. The positions of the phylogroups along these horizontal lines correspond to the percentage of sequences carrying the ancestral genes. The cryptic clade I was used as outgroup in order to confirm the intermediate evolutionary position of EPEC-503225 and KTE146 strains (see main text)

### Differences in the evolutionary pathways of the major *E. coli* root lineages

The obtained phylogenetic reconstructions suggest that three lineages were involved in the initial diversification steps. To investigate if the diversification of the lineages could be associated with particular lifestyles and evolutionary strategies, several genomic markers were analyzed such as the number of mutations per site, ancient recombination between and within phylogroups, and the gain or loss of genes.

The accumulated mutations per site revealed that, independently from the layers analyzed, the EB1A lineage presented a number of mutations below the mean value, whereas the FGB2 lineage showed values above the mean (S5 Fig). This indicates a higher mutation frequency in the FGB2 root lineage compared with the EB1A lineage. In addition, the number of accumulated mutations in the deepest branch of FGB2 lineage was lower than in recent branches, whereas in EB1A lineage the opposite occurred; more changes were accumulated in the deepest branch. The analysis of ancient recombination events revealed that around 3% of the genes belonging to *E. coli* core genome had suffered recombination events. However, the recombination frequency was not homogeneous across different phylogroups, and was more frequently found in phylogroups G and F (S6 Fig). Moreover, no ancient recombination events were detected in the phylogroup B2, belonging to FGB2 lineage. The results of accumulated mutations per site are similar without the suspected recombinant genes, as the effect of recombinatorial events is minimized when the number of genes analyzed increases [20]. Finally, to find a possible signal that might reveal the different evolutionary strategies between two root lineages in *E. coli* was the study of the gain and loss of genes. Based on the Clusters of Orthologous Groups of proteins (COG), which classify the potential products of the studied genes in functional categories, we analysed four general categories (cell interactions, replication, metabolism, and other functions). Different patterns were observed between EB1A and FGB2 lineages in these established categories (Fig 5). In general, FGB2 lineage gained more genes related to cell interaction, metabolism and replication than EB1A lineage.

**Fig 5.**
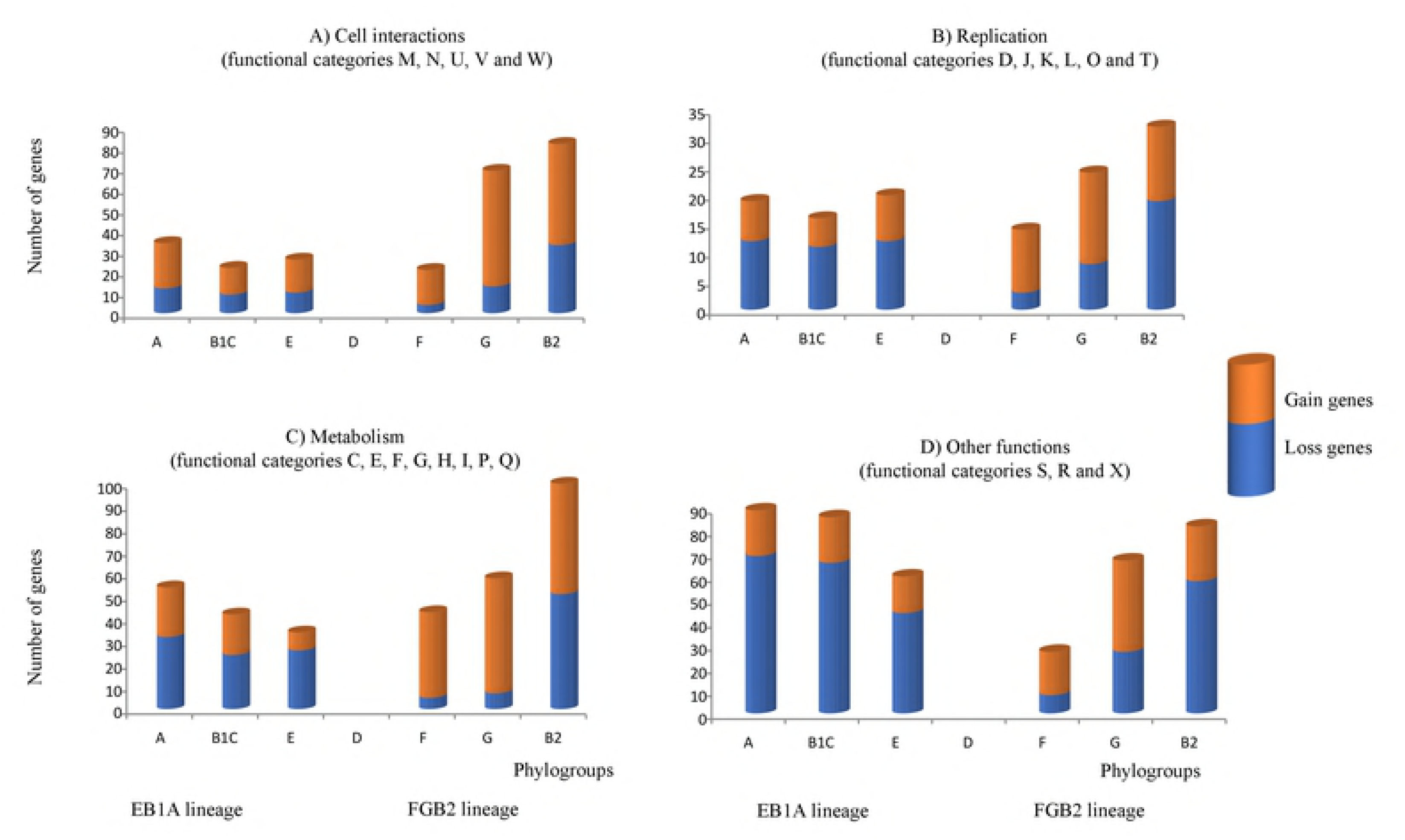
Patterns of distribution by functional categories of gained/lost gene based on Cluster Orthologous Genes (COG) classification. Four main categories were analyzed: <u>Cell interactions</u>, including the functional categories M (cell wall/membrane/ envelope biogenesis), N (cell motility), U (intracellular trafficking/ secretion/ transport), V (defense mechanism) and W (extracellular structure). <u>Replication</u>, including D (cell division), J (replication, ribosomal and biogenesis), K (transcription), L (replication, recombination and repair), O (post-translational modification, chaperones), T (signal transduction). <u>Metabolism</u>, including C (energy production), E (amino acid transport), F (nucleotide transport), G (coenzyme transport); I (lipid transport), P (inorganic ion transport), Q (secondary metabolites). <u>Other functions</u>, including S (unknown), R (general functions) and X (mobilome, prophage). Phylogroup D was used as reference genome because the number of available sequences in the previously used outgroups was very low.

### Gain and loss of genes with assigned functions involved in the different adaptive processes in the main lineages

In the last sections, we describe the evidence we found of the evolutionary differences among the main *E. coli* lineages. Our next step was investigated the acquired or eliminated genes among the members of the same phylogroup or lineage, searching for possible phylogroup-specific ecological adaptations. Obviously, the three linages should be compared with the *E. coli* ancestral genome, but this ancestral genome is no longer available (only two genomic sequences, EPEC-503225 and KTE146, could be close to the *E. coli* ancestral genome). Therefore, as the closest densely populated phylogroup to ancestral *E. coli* genome was phylogroup D, this phylogroup/lineage was used as reference in these studies. In a first analysis, several independent acquisitions with respect to phylogroup D were identified in different branches suggesting convergent evolution events. For instance, the EB1A root lineage acquired *yafQ-dinJ*, a toxin-antitoxin system, and *creBC*, a functional two-component system. This last system, involved in peptidoglycan recycling, promotes increased resistance against colicins M and E2, and is also involved in bacterial fitness and biofilm development, especially in the presence of subinhibitory β-lactam concentrations. The *yafQ-dinJ* system was also acquired by the F phylogroup, and CreBC by the GB2 sublineage (Fig 6). As to adhesins, if the EB1A lineage acquired the *yra* operon, the GB2 sublineage lost *ycg, ycb* and *sfm* operons present in the putative ancestor phylogroup D.

**Fig 6.**
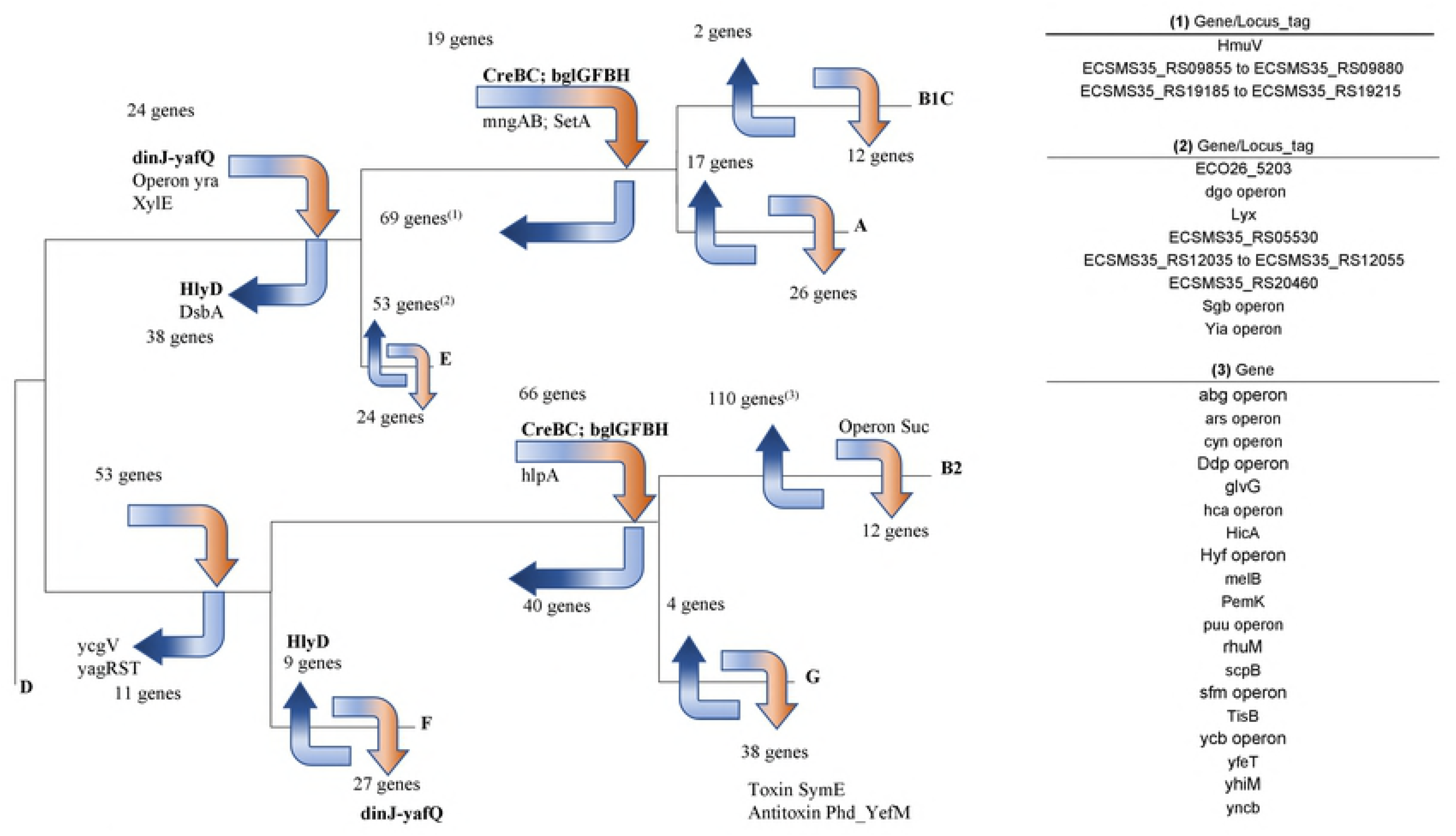
Signatures of phylogroup-core genome in the ancestral evolution of *E. coli* phylogroups. The gained/lost genes are indicated with orange and blue arrows, using the phylogeny described in this work. Genes in bold are those gained or lost in different branches indicating possible events in ecological adaptation. This representation could help to understand the different events of ecological adaptation. The genes are presented by their locus_tag identifier or with the available name in PubMed. ECSMS35 corresponds to the sequence NC_010498, ECO26 corresponds to the sequence NC_013361. Phylogroup D was used as reference genome, because the number of available sequences of the *E. coli* recent ancestor was very low.

Differences among lineages were also found with respect to genes involved in the uptake of energetic nutrients. The B1A sublineage acquired genes or operons encoding enzymes related to uptake of sugars. For instance, *bgl* operon encodes a phosphotransferase belonging to the *<u>Glc</u>*-family system is involved in the uptake of β-glucosides. Moreover, *mng* operon, belonging to the *Fru*-family, is involved in the uptake and metabolism of mannosyl-D-glycerate and *xlyE* is involved in the uptake of xylose [21]. Excess in phosphorylated sugar intermediates in B1A cells could be detrimental, causing growth inhibition [22], due to depletion of inorganic phosphate pools, which probably triggered the acquisition of the sugar efflux such as transporter encoded by *setA*, in the B1A sublineage [23]. However, the phylogroup E lost genes involved in the formation and processing of phosphorylated sugars such as xylulose 5-P, or ribose 5-P and ribulose 5-P. In addition, this phylogroup lost five genes involved in the fatty acid metabolism, suggesting deficiencies in phylogroup E for obtaining energy compared to B1A sublineage. In the FGB2 lineage, only the *bgl* operon was acquired by GB2 sublineage (Fig 6).

On the other hand, the B1A sublineage, from EB1A lineage, lost genes encoding key proteins involved in the uptake of metals as iron, manganese and molybdene, including proteins from the siderophore ABC transport system, metal-ABC transport (ECSMS35_RS09855 to ECSMS35_RS09880). Moreover, genes involved in the vitamin B12 and hemin metabolism were also lost (*hmuV*, ECSMS35_RS191855 to ECSMS35_RS19215). These genes, which might influence tissue colonization and pathogenicity, were essentially preserved in phylogroup E, suggesting that B1A sublineage could have evolved to less virulent variants compared to phylogroup E.

The FGB2 lineage lost genes involved in the detoxification of benzenic aldehydes (*yag* operon or *hca* operon) [24] and genes involved in survival in extreme conditions, such as acid pH (*hyF* operon), high temperatures and low osmolarity (*yhiM*) [25]. Moreover, the phylogroup B2 lost genes with possible environmental functions, such as transport of melobiose (*melB*), utilization of cyanate as a source of nitrogen for growth (cyn operon) or resistance to arsenate (*ars* operon). Acquisition of toxin-antitoxin related genes was found in the FGB2 lineage. In GB2 sublineage, there was a gain of *hipA* gene, which belongs to HipBA toxin/antitoxin system, and where the overexpression of the hipA protein leads to multidrug tolerance in *E. coli* [26]. In addition to the *yafQ-dinJ* system previously commented, other duplications of toxin and antitoxin genes from different systems were found in phylogroup G (*yefM* and *phD* antitoxins or *symE* toxin). The *symE* gene, encoding a toxin belonging to type I toxin-antitoxin system, has probably evolved by gene duplication [27]. Phylogroup B2 lost genes involved in toxin-antitoxin systems (*tisAB/istR, hicAB or pemI/pemK*).

## Discussion

*E. coli* is the most widely sequenced microorganism, and therefore the available material for tracing its evolutionary history is extremely abundant. However, there are discrepant aspects concerning *E. coli* phylogeny that have not been yet resolved. In this work, phylogenetic analysis of 6,220 full sequenced genomes, available in Genbank, was performed, offering some new perspectives about these open questions. During the first stages in the development of this work, some basic problems were found; for instance, around 90% of the sequenced *E. coli* genomes were not fully completed and they remain in draft [28]. If draft genomes should be or not removed from phylogenetic analysis is a matter of concern, as some genes could be lost [29]. However, the analysis of 32,000 bacterial genomes turned out the sufficient quality of the drafts for phylogenetic purposes [28]. Only six genomes in our sampling were eliminated due to poor quality of the sequences. Another observed drawback was the misallocation of 40 genomes as *E. coli* in the database. The most common mistake was to identify as *E. coli* genomes those belonging to cryptic clade I (22/40). Consequently, these misallocated genomes were excluded for the phylogenetic *E. coli* reconstruction. The cryptic clade I was used as outgroup, but the intermediate links between cryptic clade I and *E. coli* identified during this work were used as the most recent common ancestor in the analysis of staggered diversification processes in *E. coli*.

In this work, *E. coli* core genome was reduced in around 318 genes with respect to the analysis using 61 complete genomes [9]. Although it represents a drastic reduction in the number of genes, they represent around 20% of the *E. coli* genome and probably this data is coincident with previous estimations [30]. Moreover, a new phylogenetic group was identified with high support value and evolutionary divergence with respect to known phylogroups were higher than between already established phylogroups. This new phylogroup denominated as phylogroup G represented <1% of the total number of sequenced *E. coli* strains. The core genome in this new phylogroup (3,741 genes) was larger than those estimated in the previously known phylogroups (1,767-2,692 genes), but this result might be biased due to the low number of sequences belonging to this phylogroup. On the contrary, genomes initially described as phylogroup C could not be discriminated in our analysis from those of phylogroup B1. In a previous work published by our group, the phylogroup C was suspected to be composed of genomes arising by recombination between phylogroup A and B1 [16]. We could only identify members of phylogroup C as a subpopulation in phylogroup B1. On the other hand. three root lineages were defined. They were lineage D, the deepest one, EB1A and FGB2 lineages, both with three phylogenetic groups. This widely used phylogenetic reconstruction only offers the current population structure of *E. coli*, leaving the open questions, related to ancestral stepwise diversification and differentiation processes unresolved.

A new strategy for adding resolution in the evolutionary reconstruction of bacterial species combining two complementary approaches, layered phylogenomics and polymorphism fingerprinting, (LP-PF) is presented in this work. The LP approach was based on phylogenetic reconstructions with ensembles of genes corresponding to the different evolutionary stages. Those genes shared among the more separate bacterial species in the phylogenetic trees (the deepest branches) could correspond to the most ancestral genetic information (or paleome).

We examined the set of genes previously defined as minimal genome to find the ancestral traits that might cast a light about the origin of the *E. coli* species. Our interest was not to redefine the minimal genome, essentially encoding metabolic networks [31]. In fact, the number of these essential (and shared) genes used in this work (n=51) was lower than the proposed minimal set of genes in *E. coli* [32] consequently our set would be insufficient to assure the bacterial viability. We do not suggest that the minimal genome in *E. coli* could be 51 genes, we only wanted to use the highest number of genes previously identified as minimal genome present in all *E. coli* genomes for subsequent ancestral reconstructions. However, this approach showed a solid phylogenetic reconstruction but did not yield sufficient resolution for itself to infer the ancestral processes of diversification into *E. coli* phylogroups. This could be explained by the loss of ancient phylogenetic information because all available sequences in databases correspond to organisms recently sampled (last 60-70 years, mostly along the last years), and therefore, represents the phylogroups orders of magnitude as later than the first steps of the species diversification (more than 20-30 million years) [33, 34]

However, LP approach is necessary for the next step, PF approach. They are sides of the same coin. The combination of LP and PF allowed us to infer the step-by-step diversification of *E. coli* species. Single nucleotide polymorphism data in PF is now being applied to understand differentiation processes at deep evolutionary timescales since the conserved positions still maintain phylogenetic information of their ancestors (Fig 3) [35]. The results obtained using LP-PF strategy were reinforced with the analysis of the gained-lost genes in the different phylogroups with respect to hypothetical ancestral core genome (Fig 4). This evolutionary analysis strongly suggests that early steps in the diversification of *E. coli* phylogroups started with the diversification of two EB1A and FGB2 root lineages. On the other hand, the differentiation of phylogroup D only occurred much later, that is, strains from phylogroup D remained closely related with the putative common ancestor during a longer period of time, representing a different lineage. Indeed, the phylogroup D was always located in the most basal position among the known phylogroups [6, 14], conserving many traits from ancestral *E. coli* genome. Several groups had suggested a polyphyletic origin for phylogroup D [8, 13, 36]; however, one of these branches was now clearly identified as phylogroup F and the differentiation of phylogroup F was prior to phylogroup D (Fig 3). Our results also support recent studies proposing FGB2 ina different evolutionary trajectory as the first diversified root lineage [37]. In fact, phylogroup B2, was the first differentiated phylogroup and consequently the most distant with respect to origin [2, 8, 10], losing more traits than other phylogroups from the ancestral genome. Analogous results were observed in the stepwise diversification in the EB1A lineage, where A and E were the first and last that underwent differentiation into this lineage. In other words, phylogroups B2 and A represent the more evolved branches, whereas F and E the less evolved within the two root phylogroups FGB2 and EB1A, respectively.

The staggered diversification processes suggested by LP-PF strategy was used as model for new evolutionary inferences. As genomic diversification likely parallels habitat specialization, and particularly the speciation of hosts, we tried to identify any possible signal showing differences in the evolutionary strategies between these lineages. The difference in the frequency of mutations suggests a higher evolvability for the FGB2 lineage, and/or a higher ability than the EB1A lineage for the colonization of new ecological niches, and in general in transmission processes [38, 39]. On the other hand, the analysis of chromosomal sizes shows that B2 and A phylogroups (the most evolved in each respective root phylogroup) have the smallest genome sizes, whereas E and F phylogroups (the least evolved in each phylogroup) have the biggest ones (S2 Fig). These data could indicate that along the first evolutionary steps, the preservation or gain of DNA was higher than the loss, but the opposite occurred in later steps; the genetic loss was higher than the gain. This result would suggest reductive evolution processes, which had been previously proposed only for phylogroup A [1]. Moreover, when the gain and loss of specific genes was analyzed using the COG categories, the FGB2 lineage was particularly enriched in genes involved in hosts and tissues colonization, virulence with respect to the EB1A lineage, which was more endowed (particularly the B1A sub-lineage) in functions assuring a more generalist style of life (see below). These results support the concept that EB1A and FGB2 lineages could be the result of the early adoption of different adaptive strategies.

We investigated the acquired or eliminated specific functions at the time of differentiation of the different phylogenetic branches. Within the EB1A root lineage, carbohydrate transports systems (xylE, *bgl* operon and *mng* operon) required for sugars uptake (xylose, aryl beta-glucosides, and mannose respectively) were more frequently found in B1A sub-lineage. On the contrary, the phylogroup E lost genes involved in the metabolism of sugars (xylulose and ribulose) and fatty acid metabolism. This might suggest a more generalist style of life (more different available sources of energy) in the B1A sublineage. However, the acquisition of these genes involved in the sugar uptake could induce a possible detrimental effect due to an excess of phosphorylated sugars [40] in the cell, so that the B1A sublineage acquired a sugar efflux transporter (setA) to regulate the phosphorylated sugars concentration into the cell. This result is consistent with previous findings between commensal and pathogenic *E. coli* strains [41]. Similar results were also observed in the ancestral reconstruction of other microorganisms, such as enterococci, suggesting that the carbohydrates utilization has been the major driver of bacterial specialization [42]. On the other hand, the phylogroup E has genes involved in the uptake of iron in the hemin metabolism (*hmuV*, ECSMS35_RS191855 to ECSMS35_RS19215), which were missing in B1A sublineage. These genes could have contributed in the pathogenesis or in the specialization of niche and might represent an evolutionary convergence with FGB2 root lineage lifestyle (Fig 6).

Within the FGB2 root lineage, phylogroup B2, that has been suggested to be the most host-adapted including humans [43], seems to have lost some environmental-adaptive functions. These might include those involved in transport of melobiose and cyanate, or in the ability to grow in extreme conditions, such as acid pH or high temperature, or, environmentally-regulated adhesins as those encoded by *ycgV, ycb* or *sfm* (these last genes were lost by all members of the FGB2 lineage). On the contrary, EB1A root lineage acquired adhesins, as *yra*, which are only expressed as response to specific environmental changes [44].

Our analysis includes the greatest number of available whole genomes ever used to analyze the ancestral *E. coli* diversification events, offering a new and more comprehensive view on the evolutionary history of *E. coli*. Even though we used the LP-PF combined strategy to explore the ancient *E. coli* diversification events, it can be also implemented to cast light in recent diversifications, where the LP approach will probably gain more relevance. Of course, the combined strategy of LP and PF proposed in this work can be used as model for other detailed reconstructions of the evolutionary history of any other microorganism with a sufficient number of available sequenced genomes in databases. Future research on the staggered bacterial diversification will certainly provide more deep knowledge to understand the effect of environmental changes in microbial evolution.

## Materials and methods

### Data sources and selection of genes used in the different evolutionary steps

The dataset used in this work included complete and draft genome sequences of 6,290 *Escherichia* species downloaded from NCBI database (ftp.ncbi.nlm.nih.gov/genomes/genbank/bacteria/Escherichia_coli/latest_assembly_versions) as was available in August 2017. A detailed list of the genomes used is presented in S1 File. A Basic Local Alignment Search Tool (BLAST) of all-to-all genes found in the 337 complete *E. coli* genomes was performed. Genes with ≤70% similarity in amino acid sequence and ≥30% difference in sequence length were identified. This approach yielded an estimated pan-genome of 25,508 genes. Then, BLAST of each one allowed us to determine the distribution of these genes in *Escherichia* genomes included in our database.

To guarantee the correct classification of all downloaded genomes, those genes present in 100% of 6,290 available *Escherichia* genomes were defined as *Escherichia* genus core genome (S1 Table). These genes were chosen and aligned using SeaView4.4 [45]. Maximum likelihood phylogeny (ML) using GTR+ I+ Γ as a model of nucleotide substitution was estimated and visualized with SeaView program. The aLRT (approximate Likelihood Ratio Test) considered only those branches with support values > 99%. Among the genes used in *Escherichia* genus core genome were searched the genes previously identified as minimal bacterial genome [17] (S2 Table). Once the operative *E. coli* database was established, the next steps were oriented to define the species core genome, that is, the ensemble of genes present in 100% of *E. coli* genomes (S2 File).

### Framework definition

The phylogenetic reconstructions using whole genomes were based on the analysis of core and flexible genes. The core genome was defined as the set of genes present in all members belonging to the same group (normally species). The flexible genome (or accessory genome) corresponded to the set of genes that were not present in all members of the same group. The combination of core and flexible genomes among all members of a same taxonomic unit was denominated pan-genome. However, we considered that the current allocation of genes provided insufficient information to trace evolutionary trends. Trying to overcome this limitation, we applied a combined strategy based on layered phylogenomics (LP) and polymorphism fingerprinting (PF) approaches. The LP approach was based on stratifying the genes in five successive genomic subdivisions, corresponding to the minimal (essential) genome, genus-core genome and species-core genome phylogroup-core genome and phylogroup-flexible genome. Each new subdivision should carry a different set of genes giving information about the different steps in the *E. coli* evolutionary, according to Fig 1. The set of genes assigned to the minimal genome could give us the most ancestral information, as they encoded essential function for the bacterial life and consequently, they were expected to evolve from the earliest, ancestral *E. coli* times. The genus-core genome included the genes present in all members of the genus *Escherichia*, but now excluded the genes of the minimal genome, to increase the differential features in the reconstruction of the phylogroups diversification process. In a third step, the *E. coli* specie-core included the genes present in all members of *E. coli* but now excluded the genes used in the previous steps to increase the differential features in the reconstruction of the phylogroups differentiation process. Finally, remaining genes present in a phylogroup and not included in any of the core genomes were classified as phylogroup-flexible genome. They would be candidates to describe the recent events and could help us to understand the adaptive possibilities of each subpopulation into particular phylogroups and probably the future sub-specialization of subpopulations into *E. coli* phylogroups (*E. coli* expansion). The PF approach was based on the SNPs for the reconstruction at deep evolutionary timescales [35]. First, the conserved positions in all genomes of each phylogroup were known and only those with variable positions with respect to their hypothetical ancestor were selected. The number of selected SNPs was overprinted on the different branches in *E. coli* phylogeny previously established. The combined strategy could allow us to infer the evolutionary scenario of diversification in *E. coli*.

### Current and ancestral phylogeny reconstruction

To alleviate the burden of computer-time required to reconstruct large phylogenies, phylogenies of concatenated genes with cryptic clade I as outgroup (reference sequence TW10509) were reconstructed by ML with RAxML (Randomized Axelerated Maximum Likelihood) [46] using GTR + I + Γ as a model of nucleotide substitution. SH test using FastTree with values > 99% was considered valid support [47]. To classify all genomes in their corresponding phylogroups, the following reference sequences were used for the identification of the branches, NC_000913 as phylogroup A, NC_013361 as phylogroup B1, NC_009801 as phylogroup C, NC_002655 as phylogroup E, NC_017644 as phylogroup B2, NC_010498 as phylogroup F and CU928163 as phylogroup D. New monophyletic groups with more than 10 sequences were considered as new phylogroups. The orphan sequences (lower than n=10 sequences) were excluded in successive analyses. We considered as a necessary requirement to define a new phylogroup that the estimated evolutionary distance between the hypothetical new group and known phylogroups must be higher than the distance among previously established phylogroups. Evolutionary distance between two phylogroups was obtained considering the relative length of the branches. The mean intragroup evolutionary distance was estimated as the mean distance of each branch to the origin of the phylogroup, the subtree of each phylogroup was obtained from the tree and the distances were extracted with the TreeStat program included in the BEAST software (tree.bio.ed.ac.uk/software/beast/).

In order to infer the staggered diversification processes in *E. coli*, the previously described combined strategy was implemented. The phylogenetic trees in the different layers in the LP approach (minimal genome, genus-core genome and species-core genome and phylogroup-core genome) were performed using ML with RAxML using GTR + I + Γ as a model of nucleotide substitution. The SH test using FastTree with values > 99% was considered valid support. According to the PF approach, the invariant positions (100% consensus sequence) present in all genomes of the same phylogroups were identified using SeaView4. Among the conserved positions, polymorphic sites were selected using DnaSP software [48] These positions were used to reconstruct the evolutionary history using the parsimony method available in Mesquite program (www.mesquiteproject.org).

On the other hand, a second strategy based on the reconstruction of hypothetical *E. coli* ancestral core genome was implemented to reinforce the results obtained with combined LP-PF strategy. This ancestral genome was estimated by applying the MGRA program (Multiple Genome Rearrangements and Ancestors), a tool for reconstruction of ancestral gene orders and the history of genome rearrangements (mgra.cblab.org), using phylogroup-core genome and cryptic clade I as outgroup.

### Chromosomal size for all *E. coli* phylogroups

When all genomes were allocated in phylogroups, the mean chromosomal size was calculated with confidence level 95% using SPSS program. The statistical comparison among all phylogroups was estimated using Kruskal-Wallis nonparametric tests for comparing K-independent samples or the Mann-Whitney nonparametric two-sample tests.

### Inferring the accumulated mutation and recombination in each phylogroup along the time

The evolutionary distances represent the accumulated mutations per site. These data provide the mean and 95% of confidence interval of the evolutionary distances of the different ancestral branches. Those branches with values of accumulated mutations higher or lower than mean value can then be distinguished. The recombination was suspected when the topology of each gene belonging to the *E. coli*-core genome (ML using GTR + I + Γ as a model of nucleotide substitution) showed inconsistency with the topology of the *E. coli* species-core genome tree. A limitation of this approach is the lack of support for individual genes, because sometimes the phylogenetic noise is high. To avoid this limitation, the consensus phylogroup sequence for each gene (set consensus the default threshold) was defined for phylogroups. This approach reduced the noise but also excluded the non-ancestral recombination. In other words, only the ancestral recombination could be inferred. Finally, the inconsistencies were analyzed with the tree-puzzle 5.2 program [49] and SH test (p<0.05).

### Gain and lost genes between the main lineages and among different phylogroups into same lineages

This approach could identify those genes segregated during early stages of diversification/specialization. For the identification of ancestral segregation, we used a threshold of 95%-5% with respect to ancestor nodes for assigning a gene as present or absent respectively. The presence/absence of genes was inferred by parsimony method using the *E. coli* core genome phylogeny at each ancestral node, quantifying the incoming and outgoing genes between consecutive nodes of the tree. If a determined gene was lost (or gained) in two phylogroups sharing a common ancestor, only a single event (loss or gain) was considered. If they did not share a common ancestor, then we considered that two independent events had occurred. Therefore, we could then calculate how many genes and how many times the studied genes in each branch and in the *E. coli* tree were lost respectively.

Once the gain/lost genes were identified, they were classified based on their presumptive functions. The conserved domains in each gene were analyzed using CD-search tool, which allowed allocation of the genes in functional COG categories (<u>www.ncbi.nlm.nih.gov/COG</u>). We condensed these COG-categories in four supercategories: Group A: Cell interactions, including genes presumptively involved in host-bacterial interactions, including the functional COG codes M, N, U, V and W corresponding to cell wall, membrane and envelope biogenesis (M); cell motility (N); intracellular trafficking, secretion, transport (U); defense mechanism (V); extracellular structures (W). Group B: Replication, including COG codes D, J, K, L, O and T, corresponding to cell division and chromosome partitioning (D), replication, ribosomal and biogenesis (J), transcription (K), replication, recombination and repair (L), post-translational modification, protein turnover, chaperones (O), signal traduction mechanism (T). Group C: Metabolism, including C, E, F, G, H, I, P and Q, corresponding to energy production and conversion (C), aminoacid transport and metabolism (E), nucleotide transport and metabolism (F), carbohydrate transport and metabolism (G), coenzyme transport and metabolism (H), lipid transport and metabolism (I) inorganic ion transport and metabolism (P), secondary metabolites, transport and metabolism (Q). Group D: Other functions including S (unknown), R (general function) and X (mobilome, prophages and transposons). Consistent with the aim of discovering unique properties involved in the evolutionary processes of each lineage or node of diversification, the functional characteristics of genes specifically present or absent in the phylogenetic groups were examined (S3 File).

## Supporting information Legends

**S1 Fig. *Escherichia* genus phylogenetic reconstruction. A) ML tree of *Escherichia* genus-core genome**. Phylogenetic reconstruction using Maximum-Likelihood (GTR+ I+ Γ, aLR ≥99%) with the concatenate of 189 genes (244,170 nt) corresponding to 100% of sequenced *Escherichia* strains, available in Genbank (last access August-17’). All concatenate with less than 95% site coverage were eliminated. No sequence belonging to cryptic clade IV was used because there is not a complete available sequence in public database. *A. hermanni* and *E. vulneris* were also included but they were used as outgroup. **B) Distribution of those 40 misclassified *E. coli***. Identification of those sequences misclassified as *E. coli* with their access numbers (https://www.ncbi.nlm.nih.gov/nuccore/) and their correct allocation based on the previous phylogenetic reconstruction. The access numbers NZ_JNPC01000001 and NZ_JNPD01000001 have been recently re-classified as *Raoultella*.

**S2 Fig. Chromosomal size for all *E. coli* phylogroups**. The mean chromosomal size was calculated in all phylogroups with confidence range ≥95%. Significant differences in mean chromosomal sizes among phylogroups were observed (Kruskal-Wallis p<0.0001) and the pairwise comparisons were also significant (Mann-Whitney p<0.02) except for D and G phylogroups.

**S3 Fig. A) Distribution of *E. coli* genes used in the different evolutionary steps**. *E. coli* genome was differentiated in core and accessory genome. The number of genes used in the different layers are shown. The layers related to the most ancestral events (core genome) are shown in dark colors, whereas the recent events (accessory genome) are shown in high colors. The exception is the light blue oval corresponding to phylogroup-core genome as it is a core genome for phylogroups but is not E. coli core genome *senso strict*. The estimation of *E. coli* pan-genome was inferred using the 337 available complete sequences. **B) Circular maps of core genome of different phylogroups**. The inner ring corresponds to *E. coli* K2 used as reference strain. The successive rings correspond to the core genome for phylogroup A, phylogroup B1, phylogroup E, phylogroup D, phylogroup F, phylogroup G and phylogroup B2 in the most external ring.

**S4 Fig. Ancestor phylogenetic reconstruction and the origin of *E. coli* phylogroups. A) Layered phylogenomics (LP) approach**, using maximum-likelihood (GTR+I+Γ, SH ≥99%). All concatenate with less than 95% site coverage were eliminated. To avoid inferences, the genes used in the minimal genome tree were not used in *Escherichia* genus-core genome (hence although the number of genes defined as genus core genome was 189, only 138 genes were used). In a similar way, the species-core genome was performed with 838 genes, after eliminating the 189 genes corresponding to minimal and genus-core genome. The MRCA corresponds to the sequences EPEC-503225 and KTE146 identified in figure 2. A similar evolutionary reconstruction was obtained when cryptic clade I was used as outgroup, although the great evolutionary distance from *E. coli* to the cryptic clade I was confirmed by the analyzed mutations (range 76-5,545 mutations). **B) Phylogroup polymorphism fingerprinting (PF)**. Among the variable positions, only those changes present in all members of a phylogroup or lineage were analyzed. Based on the reference phylogeny Mesquite program allowed overprinting the evolutionary moment when these changes were selected. The numbers described in the parenthesis show the total number of conserved positions among all phylogroups

**S5 Fig. Inferred frequencies of accumulated mutations per site in the *E. coli* branches**. The phylogenetic reconstructions of minimal genome, genus-core genome and species-core genome were performed using ML with RAxML using GTR + I + Γ as a model of nucleotide substitution (excluding the ancestral recombinant genes). The suspected recombinant genes excluded were 5, 6 and 18 among the set of genes used in minimal genome, genus-core genome and species-core genome respectively. SH test using FastTree with values > 99% was considered valid support. The evolutionary distances represent the accumulated mutations per site. These data allow obtainment of the mean and 95% of confidence interval. The asterisks show the branches out of the confidence range. Brown and blue branches are the branches with accumulated mutations per site higher and lower than normal value respectively.

**S6 Fig. Ancestral recombination detected between the different *E. coli* phylogroups**. The recombination was suspected when the topology of the *E. coli* core genome tree showed inconsistency with the topology of the single gene tree using the consensus phylogroup sequences (set consensus default threshold). Arrows show the defection from donor to receptor.

**S1 Table. Genes defined as *Escherichia* genus core genome**. Genes present in 100% of 6,290 *Escherichia* genomes available in our database. The genes are identified by their name

**S2 Table. Genes identified as minimal bacterial genome**. Genes present in 100% of 6,290 *Escherichia* genomes available in our database and that previously has been estimated as minimal bacterial genome. The genes are identified by their name

**S1 File. Complete and draft genome sequences used in this work**. Sequences downloaded from database (<u>ftp.ncbi.nlm.nih.gov/genomes/genbank/bacteria/Escherichia_coli/latest_assembly_versions</u>). Definition includes the access number, organism and if it is a complete genome or draft

**S2 File. Genes defined as *Escherichia coli* species core genome**. Genes present in 100% of *E. coli* genomes available in our database. The genes are identified by their name or locus_tag identifier. ECNA114 corresponds to the sequence NC_017644, ECSMS35 corresponds to the sequence NC_010498, ECUMN corresponds to the sequence CU928163 and Z corresponds to the sequence NC_002655

**S3 File. Genes segregated during early stages of evolution of *Escherichia coli* phylogroups**. The presence/absence of genes were inferred by parsimony method using the *E. coli* core genome phylogeny and a stringent threshold of 95%-5% respect to ancestor nodes for assigning a gene as present or absent respectively. It includes the definition of the different COG functional categories and the reference sequences where to find the genes identified by their locus_tag.

